# T-cell hyperactivation and paralysis in severe COVID-19 infection revealed by single-cell analysis

**DOI:** 10.1101/2020.05.26.115923

**Authors:** Bahire Kalfaoglu, José Almeida-Santos, Chanidapa Adele Tye, Yorifumi Satou, Masahiro Ono

## Abstract

Severe COVID-19 patients can show respiratory failure, T-cell reduction, and cytokine release syndrome (CRS), which can be fatal in both young and aged patients and is a major concern of the pandemic. However, the pathogenetic mechanisms of CRS in COVID-19 are poorly understood. Here we show single cell-level mechanisms for T-cell dysregulation in severe SARS-CoV-2 infection, and thereby demonstrate the mechanisms underlying T-cell hyperactivation and paralysis in severe COVID-19 patients. By *in silico* sorting CD4+ T-cells from a single cell RNA-seq dataset, we found that CD4+ T-cells were highly activated and showed unique differentiation pathways in the lung of severe COVID-19 patients. Notably, those T-cells in severe COVID-19 patients highly expressed immunoregulatory receptors and CD25, whilst repressing the expression of the transcription factor FOXP3 and interestingly, both the differentiation of regulatory T-cells (Tregs) and Th17 was inhibited. Meanwhile, highly activated CD4^+^ T-cells express PD-1 alongside macrophages that express PD-1 ligands in severe patients, suggesting that PD-1-mediated immunoregulation was partially operating. Furthermore, we show that CD25^+^ hyperactivated T-cells differentiate into multiple helper T-cell lineages, showing multifaceted effector T-cells with Th1 and Th2 characteristics. Lastly, we show that CD4^+^ T-cells, particularly CD25-expressing hyperactivated T-cells, produce the protease Furin, which facilitates the viral entry of SARS-CoV-2. Collectively, CD4^+^ T-cells from severe COVID-19 patients are hyperactivated and FOXP3-mediated negative feedback mechanisms are impaired in the lung, while activated CD4^+^ T-cells continue to promote further viral infection through the production of Furin. Therefore, our study proposes a new model of T-cell hyperactivation and paralysis that drives pulmonary damage, systemic CRS and organ failure in severe COVID-19 patients.

## Introduction

Negative regulatory mechanisms of T-cell activation control inflammation in cancer, autoimmunity, and infections thus preventing excessive and prolonged inflammation which can induce tissue destruction, or *immunopathology*. Immune checkpoints such as CTLA-4 and PD-1 are well known examples of T-cell intrinsic regulators. Upon recognizing antigens, T-cells are activated and start to express PD-1 and CTLA-4, which in turn suppresses the two major signalling pathways for T-cells: T-cell receptor (TCR) signalling and CD28 co-stimulation.^1^ In addition, the transcription factor Foxp3 can be induced in activated T-cells, especially in humans, and plays a key role as an inducible negative regulator during inflammation.^2^

COVID-19 is caused by the coronavirus SARS-CoV-2, which is closely related to the severe acute respiratory syndrome coronavirus (SARS-CoV). The major symptoms such as cough and diarrhoea in mild to moderate patients can be understood through the type of tissues that can be infected by SARS-CoV-2. SARS-CoV-2 binds to angiotensin-converting enzyme 2 (ACE2) on the surface of human cells through its spike (S) protein. Viral entry is enhanced by the type II transmembrane serine protease TMPRSS2, which cleaves a part of S protein and thereby exposes the fusion domain of the S-protein.^3,4^ SARS-CoV-2 establishes infections through epithelial cells in the upper and lower airways, which express ACE2 and TMPRSS2.^5^ In addition, the pro-protein convertase Furin activates the S-protein of SARS and SARS-CoV-2.^6,7^ Intriguingly, T-cell specific deletion of *Furin* results in the impairment of effector T-cells and regulatory T-cells (Tregs) and leads to the development of age-related autoimmunity, which is accompanied by increased serum IFN-γ, IL-4, IL-6, IL-13, and IgE.^8^ In addition, Furin is preferentially expressed by Th1 cells and is critical for their IFN-γ production.^9^ As evidenced in a parasite infection model, Furin-deficient CD4^+^ T cells are skewed towards a Th2 phenotype. ^10^

It is poorly understood how SARS-CoV-2 induces severe infection in a minority of patients, who develop respiratory distress and multiorgan failure. These severe patients show elevated serum cytokines, respiratory failure, haemophagocytosis, elevated ferritin, D-dimer, and soluble CD25 (IL-2R α chain, sCD25), which are characteristic features of secondary haemophagocytic lymphohistiocytosis (sHLH)-like conditions or cytokine release syndrome (CRS). In fact, severe COVID-19 patients have elevated levels of prototypic CRS cytokines from innate immune cells including IL-6, TNF-α, and IL-10.^11,12^ Recently McGonagle et al. proposed that activated macrophages drive immune reactions that induce diffuse pulmonary intravascular coagulopathy, or *immunothrombosis*, in severe COVID-19 patients.^13^ While this may explain the unique pulmonary pathology of severe COVID-19 patients, the underlying molecular mechanisms are poorly understood.

Importantly, CRS in severe COVID-19 patients may be more systemic and involve a wide range of T-cells. Firstly, circulating T-cells are severely reduced in severe SARS-CoV-2 infections for unknown reasons ^12,14^. Intriguingly, severe COVID-19 patients show elevated serum IL-2 and soluble CD25 (IL-2 receptor α chain).^11,12^ Since IL-2 is a potent growth factor for CD25-expressing activated T-cells ^15^, the elevation of both IL-2 and CD25 indicates that a positive feedback loop for T-cell activation is established and overrunning in severe patients. These collectively highlight the roles of T-cells in the pathogenesis of severe SARS-CoV-2 infection, although the pathogenetic mechanisms are largely unknown.

In this study, we analysed the transcriptomes of CD4^+^ T-cells from a single cell RNA-seq dataset from a recent study ^16^ and thereby investigated the gene regulation dynamics during SARS-CoV-2 infection. We show that SARS-CoV-2 induces multiple activation and differentiation processes in CD4^+^ T-cells in a unique manner. We identify defects in Foxp3-mediated negative regulation, which accelerates T-cell activation and death. In addition, by analysing multiple transcriptome datasets, we propose the possibility that those abnormally activated T-cells enhance viral entry through the production of Furin in severe COVID-19 patients.

## Results

### CD4^+^ T-cells from severe COVID-19 patients highly express a unique set of activation-induced genes

We recently showed that, using scRNA-seq analysis, melanoma-infiltrating T-cells are activated *in situ* and differentiate into either FOXP3^high^ activated Tregs or PD-1^high^ T follicular helper (Tfh)-like effector T-cells ^17^. We hypothesised that those mechanisms for T-cell activation and differentiation in inflammatory sites are altered in COVID-19 patients. To address this issue, we analysed the scRNA-seq data from bronchoalveolar lavage (BAL) fluids of mild and severe COVID-19 patients.^16^

Firstly, we performed *in silico* sorting of CD4^+^ T-cells and analysed their transcriptomes (**Fig. 1a**). We applied a dimensional reduction to the CD4^+^ T-cell data using Uniform Manifold Approximation and Projection (UMAP) and Principal Component Analysis (**Supplementary Fig. 1a**). As expected, most T-cells were from either mild or severe COVID-19 patients. Notably, clusters 1, 2, 3, 5, 6, and 7 did not contain any cells from healthy controls (HC) (**Fig. 1b** **and** **1c**), indicating that these cells uniquely differentiated during the infection, regardless of whether it was mild or severe disease.

**Figure 1.**
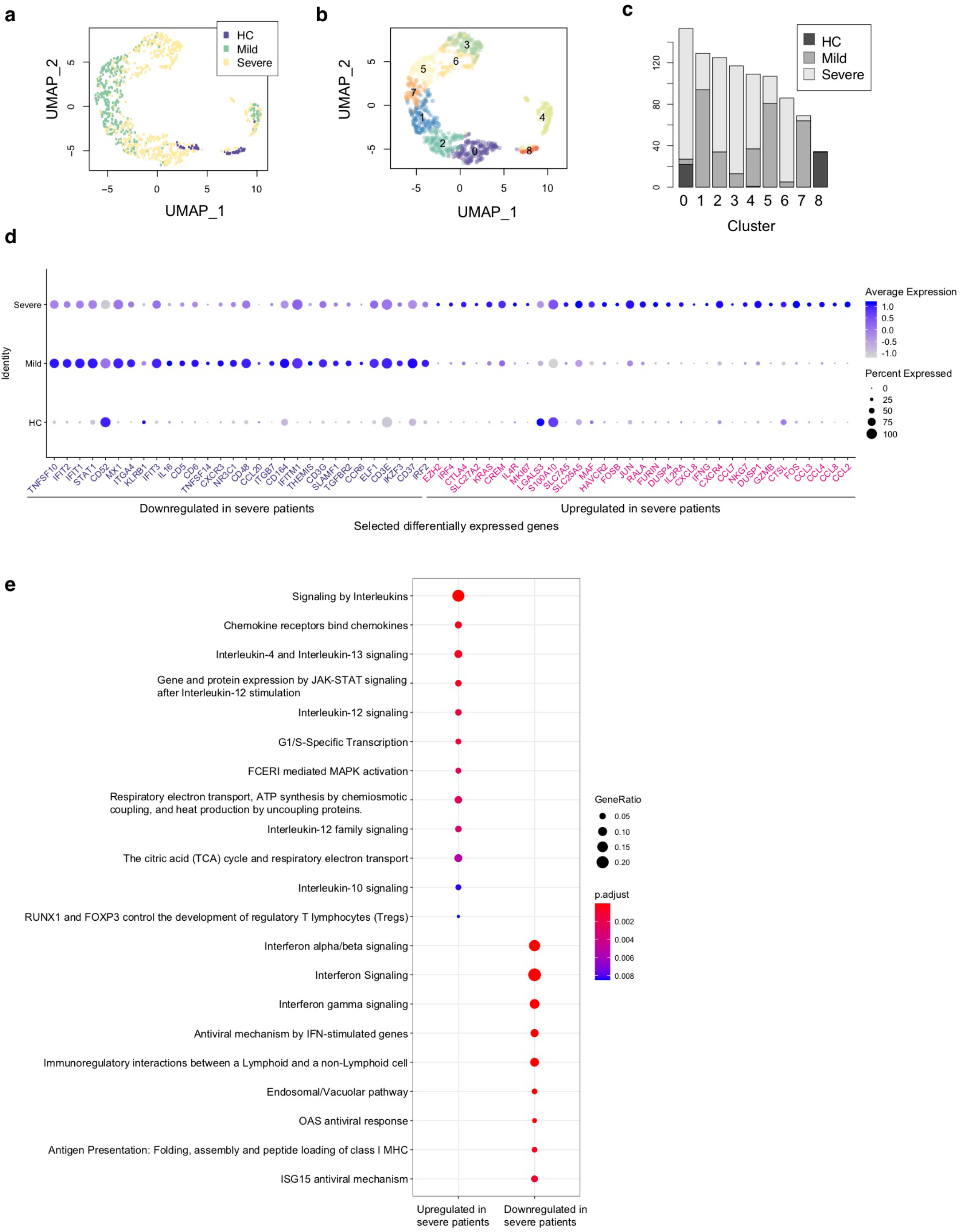
Single cell transcriptomes of CD4+ T-cells from COVID-19 patients. (a) UMAP analysis of *in silico* sorted CD4+ T-cells from COVID-19 patients. The colour code indicates the groups of patients: healthy control (HC), mild and severe COVID-19 patients. (b) Clustering of single cells in the UMAP space, showing 9 Clusters (Clusters 0 - 8). (c) The proportion of single cells from each group in each cluster. (d) Dot plots showing the expression of selected differentially expressed genes between severe and mild patients. (e) Pathway analysis of the differentially expressed genes.

Differential gene expression analysis showed that in comparison to mild patients, CD4^+^ T-cells from severe COVID-19 patients expressed higher levels of the AP-1 genes *FOS*, *FOSB*, and *JUN*, the activation marker *MKI67* (Ki67), Th2-related genes *IL4R* and *MAF*, and chemokines including *CCL2*, *CCL3*, *CCL4*, *CCL7*, *CCL8*, and *CXCL8* (**Fig. 1d, Supplementary Fig. 1b**). These suggest that CD4^+^ T-cells in severe COVID-19 patients are highly activated in the lung, recruiting macrophages, T-cells, and other immune cells. Notably, CD4^+^ T-cells in severe patients expressed higher levels of immunoregulatory genes including immune checkpoints (*CTLA4*, *HAVCR2* [TIM-3], and *LGALS3* [Galectin-3]) as well as the Tregs and T-cell activation marker *IL2RA* (CD25) (**Fig. 1d, Supplementary Fig. 1b**). These were further confirmed by pathway analysis, which identified interleukin, JAK-STAT, and MAPK signalling pathways as significantly enriched pathways (**Fig. 1e**).

On the other hand, CD4^+^ T-cells from severe patients showed decreased expression of interferon-induced genes including *IFIT1*, *IFIT2, IFIT3*, and *IFITM1* (**Fig. 1d**). Pathway analysis also showed that CD4^+^ T-cells from severe patients expressed lower levels of the genes related to interferon downstream pathways (**Fig. 1e**), suggesting that type-I interferons are suppressed in severe patients. Notably, CD4^+^ T-cells in severe patients showed lower expression of the TNF superfamily ligands *TNFSF10* (TRAIL) and *TNFSF14* (LIGHT) and the surface protein *SLAMF1* and *KLRB1*, all of which have roles in viral infections ^18–21^.

### Gene regulation dynamics for T-cell hyperactivation in severe COVID-19 patients

Next, we performed a pseudotime analysis in the UMAP space, identifying two major trajectories of T-cells, originating in Cluster 0 (**Fig. 2a, 1b**). Pseudotime 1 (*t1*) involved Clusters 0, 2, 1, 7, 5, 6, and 3, showing a longer trajectory, while pseudotime 2 (*t2*) involved Clusters 0, 8, and 4. Interestingly, the cells in the origin showed high expression of IL-7 receptor (*IL7R*), which is a marker of naïve-like T-cells in tissues.^17^ The expression of *IL7R* was gradually downregulated across the two pseudotime trajectories (t1 and t2, **Fig. 2b**). Intriguingly, T-cells at the ends of both trajectories included *MKI67* (Ki-67)^+^ T-cells and some cells were *NR4A1*^+^ or *NR4A3*^+^ (**Fig. 2c, Supplementary Fig S1c**), which indicated activation and cognate antigen signalling^22^. These analyses support that the trajectories successfully captured two major pathways for T-cells to be activated during the infection.

**Figure 2.**
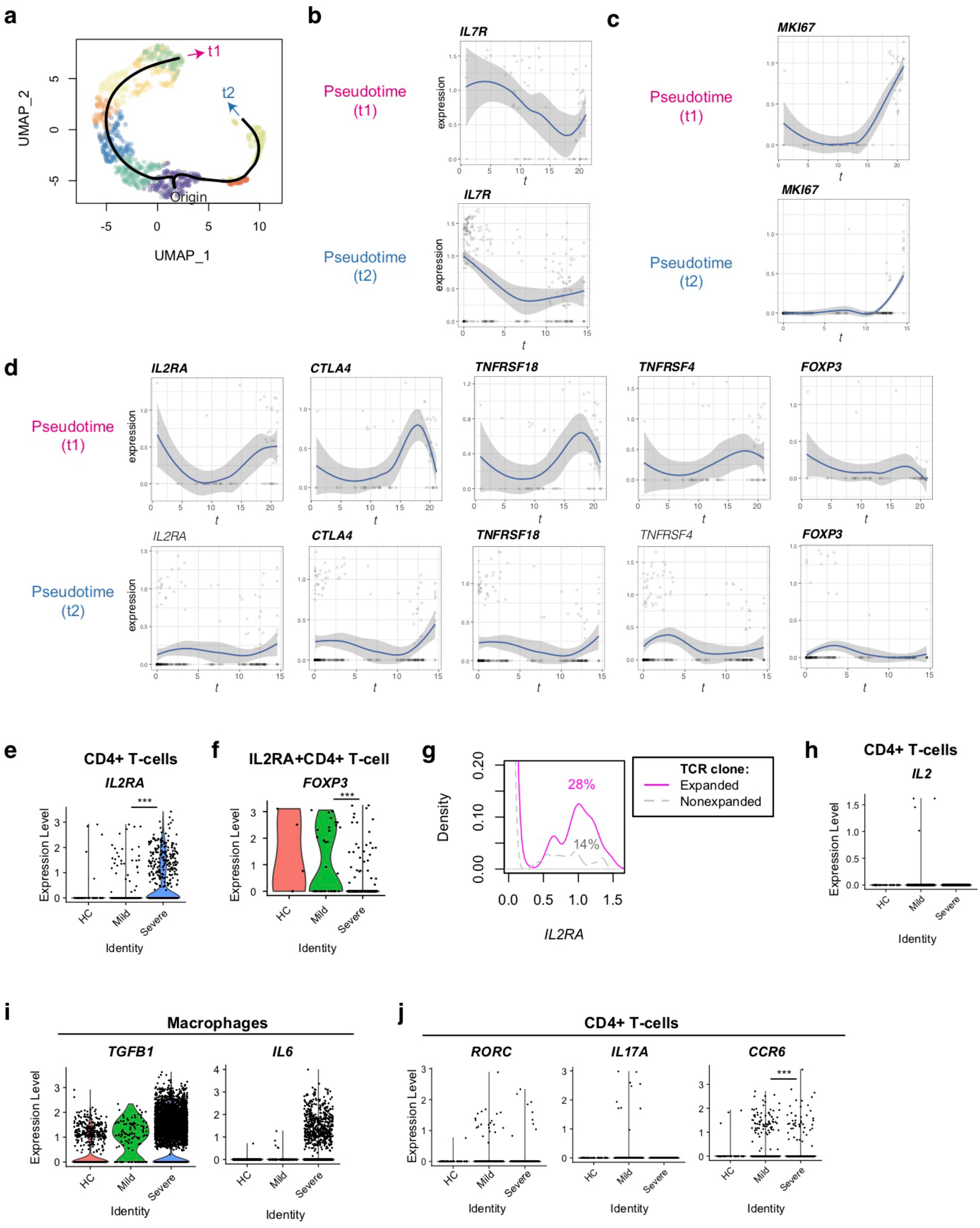
Pseudotime analysis of CD4+ T-cells from Covid-19 patients for Treg-associated genes. (a) Two pseudotime trajectories were identified in the UMAP space. (b, c) The expression of (b) *IL7R* and (c) *MKI67* in the pseudotime trajectories. (d) Gene expression dynamics of Treg-associated genes in the pseudotime trajectories. (e, f) The expression of *IL2RA* in (e) CD4+ T-cells and (f) *IL2RA*+CD4+ T-cells from the three groups. (g) *IL2RA* expression in CD4+T-cells with expanded TCR clones (n ≥ 2, magenta, solid line) and those with unexpanded TCR clones (n = 1, grey, broken line). Numbers indicate the percentage of *IL2RA*+ cells in each group. (h) The expression of *IL2* in CD4+ T-cells from the three groups. (i) The expression of *TGFB1* and *IL6* in macrophages from the three groups. (j) The expression of the Th17 genes including *RORC*, *IL17A*, and *CCR6* in CD4+ T-cells from the three groups.

Interestingly, well-known immunoregulatory genes including *IL2RA*, *CTLA4*, *TNFRSF18*, and *TNFRSF4* were more expressed in T-cells across pseudotime 1 than pseudotime 2 (**Fig. 2d**, **Supplementary Fig. 1b**). Although these genes are often associated with Tregs, *FOXP3* was not induced in both of these trajectories, and thus most of the T-cells did not become Tregs. T-cells in pseudotime 2 showed modest increase of *CTLA4* and *TNFRSF18* only towards the end of the trajectory (**Fig. 2d**).

Furthermore, *IL2RA* was significantly upregulated in CD4^+^ T-cells from severe COVID-19 patients compared to mildly affected patients (**Fig. 2e**). Since CD25 (*IL2RA*) is a key marker for Tregs and activated T-cells ^23^, we asked if CD25-expressing T-cells in COVID-19 patients were Tregs. Intriguingly, the percentage of *FOXP3*^+^ cells in *IL2RA^+^* CD4^+^ T-cells was significantly reduced in severe patients compared to mild patients (15.3% vs 48.6%, **Fig. 2f**). This indicates that *FOXP3* transcription is repressed in CD25^+^ T-cells further suggesting that *IL2RA^+^* T-cells are activated T-cells or ‘ex-Tregs’ (i.e. effector T-cells that were used to express FOXP3 but more recently downregulated FOXP3 expression^24,25^), rather than functional Tregs.

Since T-cells in the late phase of pseudotime 1 upregulated *IL2RA* and *MKI67*, we asked if expanded TCR clones expressed more *IL2RA*. Expanded TCR clones in severe patients had more *IL2RA*^+^ T-cells than those in mild patients (14% and 28% cells in mild and severe patients expressed *IL2RA*, respectively; p < 0.001, **Fig. 2g**), confirming that *IL2RA*^+^ T-cells are associated with the severe phenotype. However, no significant difference was observed between expanded and non-expanded TCR clones in severe patients. Notably, *IL2* transcription was not induced in CD4^+^ T-cells in severe patients (**Fig. 2h**), suggesting that CD25^+^ activated T-cells in severe patients die, at least partly, by cytokine deprivation.

Given that T-cells from severe patients dominated in the last part of pseudotime 1 (i.e. Clusters 3 and 6), these findings indicate that T-cells become more activated and vigorously proliferate in severe COVID-19 patients than mild patients. These CD25^+^ activated T-cells are likely to be short-lived and do not initiate *FOXP3* transcription in severe COVID-19 patients, while they can differentiate into Tregs in moderate infections.

### The differentiation of Th17 is suppressed in severe COVID-19 patients despite IL-6 availability

Firstly, we hypothesized that *FOXP3* transcription was actively repressed by cytokines in the microenvironments in severe COVID-19 patients. *FOXP3* transcription is activated by IL-2 and TGF-β signalling and is repressed by IL-6 and IL-12 signalling.^26^ In fact, some macrophages from severe COVID-19 patients expressed *TGFB1* and *IL6* (**Fig. 2i**), as shown by Liao et al.^16^ However, CD4^+^ T-cells did not increase Th17-associated genes including *RORC*, *IL17A*, and *IL17F*, and the expression of CCR6, a marker for Th17 cells, was significantly reduced in severe COVID-19 patients (**Fig. 2j** and **Supplementary Fig. 1d**). This suggests that the differentiation of both Tregs and Th17 is inhibited.

Th17 differentiating T-cells express both Foxp3 and RORγ-t before they mature.^27^ In addition, FOXP3^intermediate^ CD45RA^-^ T-cells express RORγ-t and Th17 cytokines.^28^ Together with the scRNA-seq analysis results above, these support the model that activated T-cells show differentiation arrest or preferentially die before becoming Tregs or Th17 cells in severe COVID-19 patients. Importantly, *IL2RA* expression was significantly increased in severe COVID-19 patients in comparison to mild patients, whilst very few T-cells expressed *IL2* (**Fig. 2h**).

### Lack of Tfh differentiation and evidence of PD-1-mediated immunoregulation

PD-1 is another key immunoregulatory molecule for suppressing immune responses during viral infection.^29^ However, PD-1 may play multiple roles in CD4^+^ T-cells, as PD-1 is a marker for Tfh. In fact, PD-1^high^ BCL6^high^ Tfh-like T-cells are a major effector population in melanoma tissues.^17^ Thus we asked if PD-1-expressing T-cells show Tfh differentiation and/or if PD-1-expressing T-cells can succumb to PD-1 ligand-mediated inhibition in COVID-19 patients. However, in SARS-CoV-2 infection, *BCL6* was not induced in the major activation and differentiation pathway, pseudotime 1, indicating that those activated T-cells did not differentiate into Tfh. Comparatively, T-cells in pseudotime 2 showed some upregulation, although this was statistically not significant (**Fig. 3a**). This suggests that Tfh differentiation was suppressed in COVID-19 patients. *PDCD1* was highly upregulated in both pseudotime 1 and 2, suggesting that these cells are vulnerable to PD-1 ligand-mediated suppression. Interestingly, macrophages from severe COVID-19 patients expressed higher levels of *CD274* (PD-L1) yet the expression of *PDCD1LG2* (PD-L2) was not significantly different between mild and severe patients (**Fig. 3b**). These results indicate that PD-1-mediated T-cell regulation was at least partially operating in severe COVID-19 patients. Meanwhile, *TBX21* and *GATA3* expression is induced in T-cells across pseudotime 1, suggesting that these T-cells may differentiate into Th1 and Th2 cells (**Fig. 3c**).

**Figure 3.**
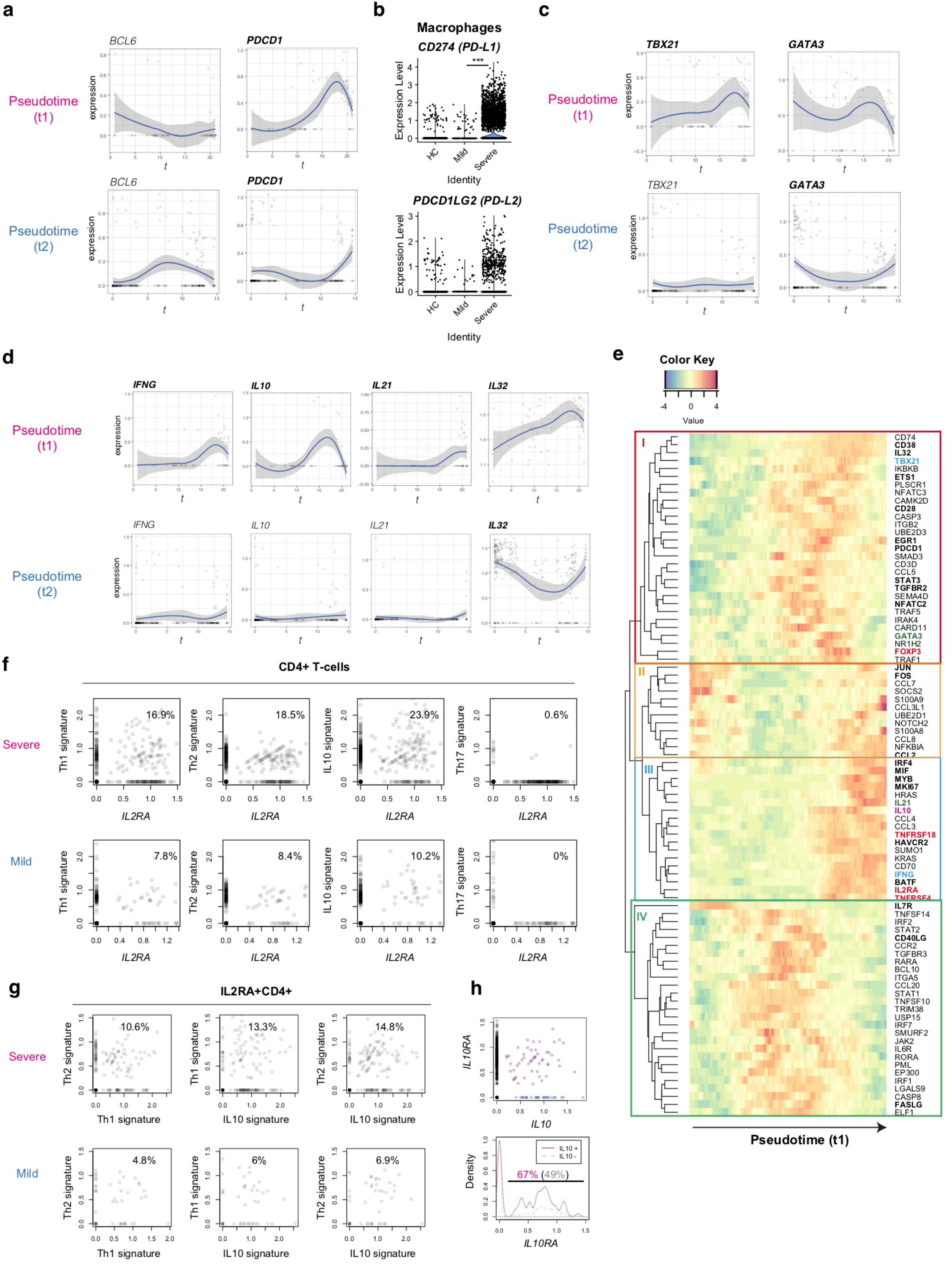
Analysis of the expression dynamics of effector T-cell genes in CD4+ T-cells from COVID-19 patients. (a) The expression of *BCL6* and *PDCD1* in the pseudotime trajectories. (b) The expression of PD1 ligand genes, PD-L1 (*CD274*) and PD-L2 (*PDCDLG2*), in macrophages from the three groups. (c) The expression of Th1 transcription factor, *TBX21*, and Th2 transcription factor, *GATA3*, in the pseudotime trajectories. (d) Gene expression dynamics of *IFNG*, *IL10*, *IL21* and *IL32* in the pseudotime trajectories. (e) Heatmap analysis of selected genes in pseudotime 1 (t1). Here the rows are key genes that are differentially expressed across pseudotime 1, and the columns are single cells in the order of pseudotime 1. Th1, Th2, and Treg-associated genes are highlighted by cyan, green, and red. In addition, key genes are highlighted as bold text. (f, g) The expression of signature genes in (f) CD4+ T-cells and (g) *IL2RA*+CD4+ T-cells. (h) (upper panel) The expression of *IL10* and *IL10RA* in CD4+ T-cells from COVID-19 patients. *IL10*+*IL10RA*+ double positive cells are highlighted by purple, while *IL10*+ single positive cells are shown by blue. (lower panel) Density plots of *IL10RA* expression in *IL10*+CD4+ T-cells and *IL10*-CD4+ T-cells.

### Dissecting activation and differentiation processes in CD4^+^ T-cells from COVID-19 patients by analysing gene expression dynamics in the pseudotime trajectory

T-cells in the late phase of pseudotime 1 upregulated the expression of cytokines including *IFNG* and *IL10*, which are Th1 and Th2 cytokines, respectively (**Fig. 3d**). In addition, *IL21* (a Th2 and Th17 cytokine) was upregulated in some cells in pseudotime 1 whereas IL-32 was highly sustained in both of the pseudotime trajectories. These indicate that differentiation processes for T-cell lineages are simultaneously induced in activated T-cells from the lung of COVID-19 patients.

Accordingly, we hypothesized that CD25-expressing activated T-cells preferentially differentiate into effector T-cells in severe COVID-19 patients, instead of their most frequent fate as Tregs in a normal setting.^23^ In order to test this hypothesis and reveal dynamics of each T-cell differentiation pathway, we analysed co-regulated genes across pseudotime 1, obtaining 4 gene modules by a hierarchical clustering (**Fig. 3e**).

Heatmap analysis of pseudotime 1 successfully captured the pseudo-temporal order of gene expression: genes in *modules II* and *IL7R* are firstly activated, followed by genes in *module IV* (apart from *IL7R*), subsequently by genes in *module I*, and lastly genes in *module III* alongside *module II* genes again (**Fig. 3e**). Reasonably, *module II* contained the AP-1 transcription factors *FOS* and *JUN,* suggesting that T-cells that highly express these genes have been recently activated. In pseudotime 1, these *FOS^+^JUN^+^* T-cells were followed by T-cells with high expression of genes in *module IV*, which contained *CD40LG* and *FASLG* (**Fig. 3e**). These *CD40LG^+^FASLG^+^* T-cells are considered to activate CD40-expressing macrophages and dendritic cells as well as inducing apoptosis of FAS-expressing cells by providing CD40 signalling and Fas signalling upon contact.

*CD40LG^+^FASLG^+^* T-cells are followed by T-cells that highly expressed genes in *module I*, which include the Th1 transcription factor *TBX21*, the Th2 transcription factor *GATA3*, and *FOXP3*. In addition, these T-cells upregulated the immediate early genes *EGR1* and *NFATC2* and the activation markers *CD38* and *PDCD1* (**Fig. 3e**). These collectively suggest that those *CD38^+^ PDCD1^+^* T-cells received sustained TCR signals and became activated, expressing *TBX21*, *GATA3*, and *FOXP3* as TCR signal downstream genes.

Lastly, T-cells in the final part of pseudotime 1 upregulated the expression of genes in *modules II* and *III*. *Module III* contained the cytokines *IL10* and *IL21*, immune checkpoints *TNFRSF4* (OX-40) and *HAVCR2* (TIM-3), Tregs and activation marker *IL2RA* and *TNFRSF18* (GITR), all of which were found to be upregulated (**Fig. 3e**). On the other hand, *FOXP3*, *TBX21* and *GATA3* were largely repressed in the T-cells. Intriguingly, those T-cells highly expressed the transcription factors for the differentiation of effector Tregs (i.e. activated Tregs with enhanced immunoregulatory activities) including *IRF4*, *MYB*, and *BATF*. Therefore, those activated T-cells partially show an effector Treg phenotype but lack their cardinal immunoregulatory features because *FOXP3* expression is repressed and the effector cytokines are transcribed. Notably, those activated T-cells also upregulated AP-1 (*FOS*/*JUN*) and other genes in *module II*, suggesting that these CD25^+^ activated T-cells had received TCR and/or costimulatory signals such as TNFRSF signals. Given that this last phase of pseudotime 1 (i.e. UMAP clusters 3 and 6) is dominated by CD4^+^ T-cells from severe patients, these results collectively support that FOXP3-mediated negative feedback on T-cell activation is defective in COVID-19 patients.

### Evidence of the differentiation of CD25-expressing activated T-cells into multifaceted effector T-cells

In order to further understand why CD25-expressing T-cells failed to differentiate into effector Tregs, we hypothesized that CD25-expressing activated T-cells are more likely to differentiate into multiple effector T-cell subsets in severe COVID-19 patients than mildly affected individuals. In fact, *IL2RA*^+^CD4^+^ T-cells from severe COVID-19 patients expressed Th1, Th2, and IL-10 signature genes more frequently (**Fig. 3f**) whereas Th17 differentiation was suppressed in *IL2RA^+^* T-cells from both groups.

Although IL-10 has been classically regarded as a Th2 cytokine, Th1 cells can produce IL-10.^30^ In fact, *IL4, IL5, IL12A, and IL13,* were not detected in any T-cells analysed in the dataset (data not shown). Thus, we asked if Th2 differentiation was diverted into IL-10 producing immunoregulatory T-cells (Tr1), which differentiate by IL-10 signalling and produce IL-10 and thereby suppress immune responses particularly in mucosal tissues.^31^. Surprisingly, however, higher frequencies of *IL2RA^+^* T-cells expressed Th1 and Th2 signature genes, concomitantly expressing IL10 signature genes (**Fig. 3g**). This strongly supports that IL-10 producing T-cells are not immunoregulatory but Th1-Th2 multifaceted effector T-cells.

*IL10RA* and *IL10RB* were expressed in activated T-cells in both of the trajectories (**Supplementary Fig. 1c**), and the frequency of *IL10RA*^+^ cells was significantly higher in *IL10*^+^ cells than *IL10*-cells (67% vs 49%, p = 0.003, **Fig. 3h**). This suggests that a positive feedback loop for IL-10 expression promoted the differentiation of the multifaceted effector T-cells in an autocrine manner ^32,33^.

### FURIN is induced in activated CD4^+^ T-cells upon TCR signals and during COVID-19 infection

Intriguingly, *FURIN* expression was increased in CD4^+^ T-cells from severe COVID-19 patients (**Fig. 1d**). Furin was previously associated with Treg functions in a knockout study^8^, although the underlying mechanisms were not clear. In addition, it was not known if Furin was specifically expressed in Tregs amongst CD4^+^ T-cells. Recently we showed that the majority of Treg-type genes are in fact regulated by TCR signalling ^17,34^, since Tregs receive infrequent-yet-regular TCR signals *in vivo* ^22^. We hypothesise that T-cells produce Furin upon activation, which can enhance the viral entry of SARS-CoV-2 into lung epithelial cells during inflammation.

Firstly, we analysed the microarray data of various CD4^+^ T-cell populations from mice ^35^. In line with our previous observations ^17^, all the antigen-experienced and activated T-cell populations including Tregs, memory-phenotype T-cells, and tissue-infiltrating effector T-cells showed higher expression of *FURIN* than naïve T-cell populations (**Fig. 4a**).

**Figure 4.**
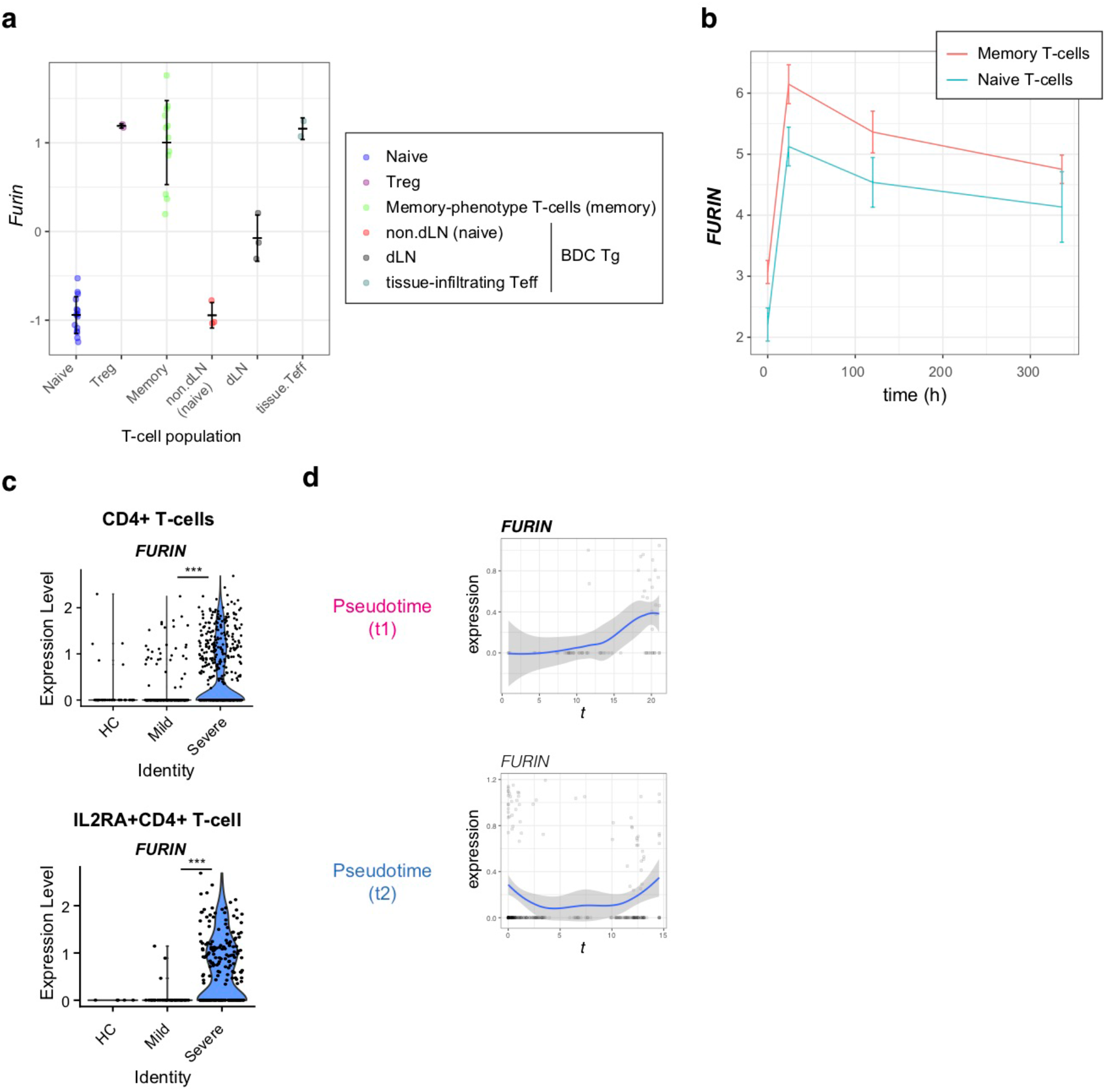
*FURIN* expression in activated CD4+ T-cells in normal conditions and COVID-19 infection. (a) *Furin* expression in CD4+ T-cell subpopulations from normal mice: naïve T-cells (naïve), Tregs, memory-phenotype T-cells (memory) from WT mice; non-draining lymph nodes (non-dLN, naïve), draining lymph nodes (dLN) of the pancreas, and pancreas-infiltrating effector T-cells (tissue-infiltrating Teff) of diabetes-prone BDC transgenic (Tg) mice. (b) Time course analysis of *FURIN* expression in human memory T-cells and naïve T-cells. (c) The expression of *FURIN* in CD4+ T-cells (upper) and *IL2RA*+CD4+ T-cells (lower) from the groups of patients and HC. (d) Gene expression dynamics of *FURIN* in the pseudotime trajectories 1 and 2.

Next, we asked if human naïve and memory CD4^+^ T cells can express *FURIN* upon receiving TCR signals. We addressed this question using the time course RNA-seq analysis of CD45RA^+^CD45RO^−^CD4^+^ naïve T-cells and CD45RA^−^CD45RO^+^ CD4^+^ memory T-cells which were obtained from 4 individuals and activated by anti-CD3 and anti-CD28 antibodies ^36^. *FURIN* expression was markedly induced within 24 hours after stimulation and was sustained over 2 weeks in the culture (**Fig. 4b**). Memory T-cells also showed higher expression of *FURIN* over the time course (p = 0.059), with a significant difference at its peak (24 hours, p = 0.004). Thus, *FURIN* expression is induced by TCR signals in human and mouse T-cells.

In SARS-CoV-2 infection, 36% of CD4^+^ T-cells and 56% of IL2RA^+^CD4^+^ T-cells from severe patients expressed *FURIN*, while in mild patients only 11% and 30% of those cells, respectively, expressed *FURIN* (**Fig. 4c**). Importantly, *FURIN* was significantly induced in CD4^+^ T-cells in pseudotime 1, particularly when T-cells upregulated CD25, CTLA-4, and TNFRSF molecules, but not in pseudotime 2 (**Fig. 4d**). These collectively support that *FURIN* expression is induced in highly activated non-regulatory CD25^+^CD4^+^ T-cells in severe COVID-19 patients.

## Discussion

Our study has shown that CD4^+^ T-cells in severe COVID-19 patients have dysregulated activation and differentiation mechanisms. The most remarkable defect was the decoupling of Treg-type activation and FOXP3 expression, which normally occurs simultaneously to sustain the effector Treg population while inflammation is resolved. ^34^

FOXP3 is induced in activated T-cells by TCR signals and its transcription is further enhanced by IL-2 and TGF-β signalling. Once expressed in T-cells, FOXP3 proteins bind to pre-existing transcription factors, particularly RUNX1 and ETS1, and thereby convert RUNX1/EST1-containing transcriptional machineries for T-cell activation and effector function into immunoregulatory ones.^34,37^ The Treg-type transcriptional regulation is characterized by the activation of immunoregulatory proteins, including many of the surface immune checkpoints such as CTLA-4, GITR, and OX-40, while cytokine genes are generally repressed. This Treg-type setting is further enhanced upon activation, when Tregs begin to show the effector Treg phenotype, further upregulating the expression of the immune checkpoint molecules. Importantly, Tregs need to sustain FOXP3 transcription in a persistent manner across time,^38^ otherwise they can downregulate FOXP3 expression and become effector T-cells.^39,40^

Since IL-2 signalling enhances *FOXP3* transcription, CD25^+^ T-cells are likely to differentiate into FOXP3^+^ Tregs in normal situations.^23^ However, in severe COVID-19 patients, those CD25^+^ T-cells are considered to be vigorously proliferating, whilst becoming multifaceted effector T-cells or dying, instead of maturing into FOXP3^+^ Tregs. Accordingly, we propose to define the unique activation status of CD25^+^FOXP3-T-cells as *hyperactivated T-cells* (**Fig. 5**). CD25 expression occurs mostly in CD4^+^ T-cells, and therefore, these CD25^+^ hyperactivated T-cells are likely to be the source of the elevated serum soluble CD25 in severe COVID-19 patients. These hyperactivated CD25^+^ T-cells produce Furin, which can further enhance SARS-CoV-2 viral entry into pulmonary epithelial cells.

**Figure 5.**
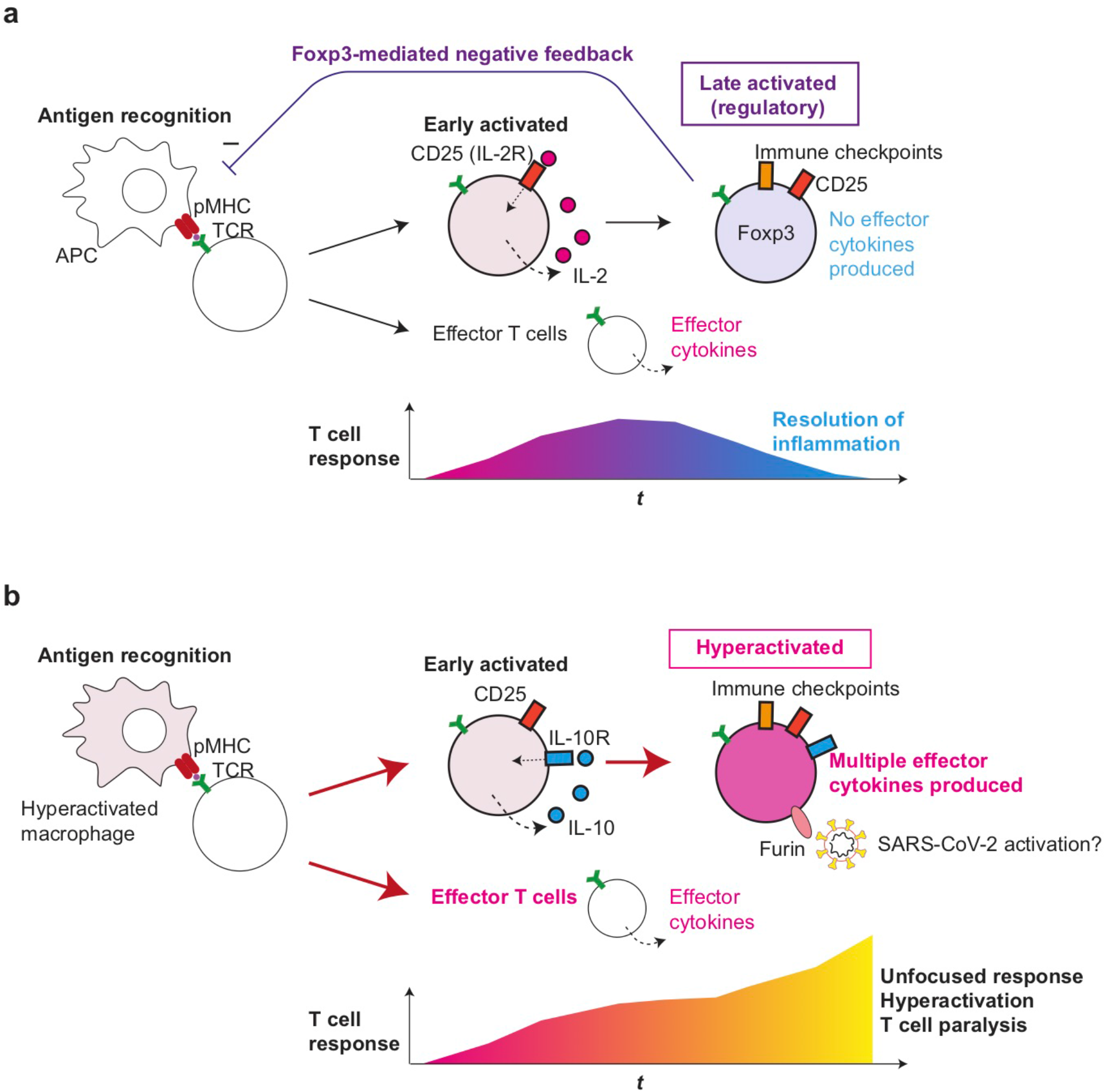
Roles of T-cell hyperactivation in the lung of severe COVID-19 patients. (a) Peripheral Treg differentiation in normal conditions. Antigen-presenting cells (APC) present antigens as peptide-MHC complex (pMHC) to CD4+ T-cells, which triggers TCR signalling and subsequent activation and differentiation processes. Initially, early activated T-cells start to produce CD25 and IL-2, establishing a positive feedback loop for T-cell activation and proliferation. Some T-cells can differentiate into effector T-cells such as Th1 and Th2. Since IL-2 signalling enhances FOXP3 transcription, prolonged activation results in the expression of immune checkpoints such as CTLA-4 and FOXP3, which represses the transcription of effector cytokine genes. CD25+CTLA-4+FOXP3+ T-cells can consume and occupy immunological resources including IL-2 and CD28 signalling, and thereby mediate a negative feedback loop on the initial T-cell activation.^23^ This leads to the suppression of T-cell responses and the resolution of inflammation. In severe COVID-19 patients, hyperactivated macrophages^13^ may present antigens to CD4+ T-cells, which are activated and differentiate into CD25+ IL10R+ early activated T-cells which produce IL-10 rather than IL-2. *FOXP3* transcription remains to be suppressed due to this and other unidentified mechanisms such as metabolism, while cytokines such as IL-10 further enhance the activation of CD25+ T-cells, resulting in the generation of CD25+ hyperactivated T-cells that express immune checkpoints, multiple effector T-cell cytokines, and Furin. The multifaceted Th differentiation may lead to unfocused T-cell responses and thereby paralyse the T-cell system. In addition, Furin can activate the S-protein of SARS-CoV-2 and thereby enhance viral entry into lung epithelial cells.

The risk factors for the development of severe COVID-19 include age, obesity, cardiovascular diseases, diabetes, and the use of corticosteroids.^41,42^ These diseases are associated with dysregulated hormonal and metabolic environments that can dysregulate the homeostasis of CD25^+^ T-cells and FOXP3-expressing Tregs. Thus, it is imperative to investigate if genes and metabolites associated with the disease conditions have any roles in promoting the differentiation of hyperactivated T-cells. Previous reports showed that Tregs accumulated in atherosclerotic lesions,^43^ and *FOXP3* expression was reduced in CD25^+^CD4^+^ T-cells from patients with prior myocardial infarctions.^44^ In addition, T-cells in patients with obesity may show different responses to T-cell activation. Intriguingly, leptin a key hormone produced by adipose tissue, is thought to prevent CD25^+^CD4^+^ T-cell proliferation but is relatively deficient in obese patients.^45,46^ Furthermore, the function of Tregs is impaired in type-1 diabetes patients.^47^ In addition, *FOXP3* transcription is transiently activated in T-cells of severe COVID-19 patients but may be repressed due to their unique metabolic states. T-cell activation is dependent on glycolysis, which converts glucose to pyruvate, and the tricarboxylic acid (TCA) cycle, which activates oxidative phosphorylation (OXPHOS) and generates ATP in mitochondria.^48^ Treg differentiation is more dependent on OXPHOS and can be inhibited by glycolysis.^49^ Importantly, our pathway analysis suggested that these metabolic pathways were altered in severe COVID-19 patients, although further studies on metabolism are required. Furthermore, the hypoxic environment in the lung of severe COVID-19 patients may activate HIF-1α, which mediates aerobic glycolysis, and thereby promotes the degradation of FOXP3 proteins.^50^ The reduction of FOXP3 proteins may result in the abrogation of the FOXP3 autoregulatory transcriptional loop thus blocking Treg differentiation.^38^

CD25^+^ hyperactivated T-cells also expressed PD-1, and PD-L1 expression in macrophages was increased in severe COVID-19 patients. This clearly shows that the PD-1 system is not able to control hyperactivated T-cells. This may be due to the status of macrophages and other antigen-presenting cells because CD80 on these cells disrupts the PD-1 - PD-L1 interaction and thereby abrogates PD-1-mediated suppression.^51^ In addition, PD-L1 expression on lung epithelial cells may play a role in regulating PD-1-expressing T-cells, as shown in other viruses including Influenza Virus and Respiratory Syncytial Virus.^52,53^

Hyperactivated T-cells differentiated into multifaceted Th1-Th2 cells with IL-10 expression. While IL-10 may serve as a growth factor for these cells through their IL-10 receptors, other cytokines in the microenvironment may drive the expression of both Th1 and Th2 transcription factors. Prototypic cytokines for Th1 and Th2 were not differentially expressed in all single cells between mild and severe patients in the current dataset. Importantly, although T-bet and Gata3 are usually considered as Th1 and Th2 transcription factors, respectively, the expression of both T-bet and Gata3 is induced in CD4^+^ T-cells by TCR signals at the early stage of T-cell activation.^54,55^ This suggests that the multifaceted Th1-Th2 T-cells are in fact still at the early stage of differentiation. Importantly, Foxp3-deficient T-cells show a similar phenotype with multifaceted Th1-Th2 differentiation. Using bacterial artificial chromosome (BAC) Foxp3-GFP reporter and Foxp3-deficient Scurfy mice, Kuczma et al showed that *Foxp3*-transcribing T-cells without functional Foxp3 proteins produced both IFN-γ and IL-4.^56^ Although the biological significance of the multifaceted Th differentiation in severe SARS-CoV-2 infection is not yet clear, we suggest that such unfocused T-cell responses will lead to the activation of broad-range of immune cells in an unorganized manner contributing to the hyperactivation as well as paralysis of the immune system in severe COVID-19 patients (**Fig. 5**).

In conclusion, our study demonstrates that SARS-CoV-2 drives hyperactivation of CD4+ T-cells and immune paralysis to promote the pathogenesis of disease and thus life-threatening symptoms in severely affected individuals. Therefore, therapeutic approaches to inhibit T-cell hyperactivation and paralysis may need to be developed for severe COVID-19 patients

## Materials and Methods

### Datasets

The single-cell-RNA-seq data from COVID-19 patients and healthy individuals was obtained from GSE145926.^16^ The microarray data of murine T-cell subpopulation were from the Immunological Genome Project (GSE15907^35^). The RNA-seq data in GSE73213^36^ was used for time course analysis of naïve and memory CD4^+^ T-cells.

### In silico sorting of CD4 T-cells

We used h5 files of the scRNA-seq dataset (GSE145926^16^) which were aligned to the human genome (GRCh38) using Cell Ranger, by importing them into the CRAN package Seurat 3.0.^57^ Single cells with high mitochondrial gene expression (higher than 5%) were excluded from further analyses. *In silico* sorting of CD4^+^ T-cells was performed by identifying them as the single cells *CD4* and *CD3E,* because no other methods, including the Bioconductor package *singleR*, reliably identified CD4^+^ T-cells. The TCR-seq data of GSE145926^16^ was used to validate the *in silico* sorting and also for analysing gene expression in expanded clones. Macrophages were similarly identified by the *ITGAM* expression and lack of *PAX5*, *CD19* and *CD3E* expressions.

### Dimensional reduction and differential gene expression

PCA was applied on the scaled data followed by a K-nearest neighbor clustering in the PCA space. UMAP was performed on clustered data using the first PCA axes. Differentially expressed genes were identified by adjusted p-values < 0.05 using the function FindMarkers of Seurat. Th1, Th2, and IL-10 signature were defined as the sum of the normalized gene expression of *IFNG*, *TBX21*, *IL21A* (Th1); *GATA3*, *IL4*, *IL6* (Th2); and *RORC*, *IL17A*, *IL17F* (Th17), respectively.

### Pathway analysis

The enrichment of biological pathways in the gene lists was tested by the Bioconductor package clusterProfiler,^58^ using the Reactome database through the Bioconductor package ReactomePA, and pathways with false discovery rate < 0.01 and q-value < 0.1 were considered significant.

### Pseudotime analysis

Trajectories were identified using the Bioconductor package *slingshot*, assuming that the cluster that show the highest expression of *IL7R* and *CCR7* is the origin. The CRAN package *ggplot2* was used to apply a generalised additive model of the CRAN package *gam* to each gene expression data. Genes that were differentially expressed across pseudotime was obtained by applying the generalised additive model to the dataset using *gam*, performing ANOVA for nonparametric effects and thereby testing if each gene expression is significantly changed across each pseudotime (p-value < 0.05).

### Other Statistical Analysis

The enrichment of cytokine-expressing single cell T-cells was tested using a chi-square test. The time course data of *FURIN* expression was analysed by one-way ANOVA with Tukey’s honest significant difference test.

## Acknowledgements

MO is supported by MRC project grant (MR/S000208/1) and is currently a Visiting Associate Professor in IRCMS, Kumamoto University to conduct the Kakenhi project 19H05426. This research was supported in part by grants from the Japan Agency for Medical Research and Development (grant numbers JP20jm0210074 to YS and MO; JP20fk0410023 and JP19fm0208012 to YS). The authors declare that no conflicts of interest exist in relation to this manuscript.

## References

1 Okazaki, T. & Okazaki, I. M. Stimulatory and Inhibitory Co-signals in Autoimmunity. Adv Exp Med Biol 1189, 213–232, doi:10.1007/978-981-32-9717-3_8 (2019).

2 Bending, D. & Ono, M. From stability to dynamics: understanding molecular mechanisms of regulatory T cells through Foxp3 transcriptional dynamics. Clin Exp Immunol, doi:10.1111/cei.13194 (2018).

3 Hoffmann, M. et al. SARS-CoV-2 Cell Entry Depends on ACE2 and TMPRSS2 and Is Blocked by a Clinically Proven Protease Inhibitor. Cell 181, 271–280.e278, doi:10.1016/j.cell.2020.02.052 (2020).

4 Glowacka, I. et al. Evidence that TMPRSS2 Activates the Severe Acute Respiratory Syndrome Coronavirus Spike Protein for Membrane Fusion and Reduces Viral Control by the Humoral Immune Response. Journal of Virology 85, 4122–4134, doi:10.1128/jvi.02232-10 (2011).

5 Sungnak, W. et al. SARS-CoV-2 entry factors are highly expressed in nasal epithelial cells together with innate immune genes. Nature Medicine, doi:10.1038/s41591-020-0868-6 (2020).

6 Coutard, B. et al. The spike glycoprotein of the new coronavirus 2019-nCoV contains a furin-like cleavage site absent in CoV of the same clade. Antiviral Research 176, 104742, doi:https://doi.org/10.1016/j.antiviral.2020.104742 (2020).

7 Hoffmann, M., Kleine-Weber, H. & Pöhlmann, S. A Multibasic Cleavage Site in the Spike Protein of SARS-CoV-2 Is Essential for Infection of Human Lung Cells. Molecular Cell 78, 779–784.e775, doi:https://doi.org/10.1016/j.molcel.2020.04.022 (2020).

8 Pesu, M. et al. T-cell-expressed proprotein convertase furin is essential for maintenance of peripheral immune tolerance. Nature 455, 246–250, doi:10.1038/nature07210 (2008).

9 Pesu, M., Muul, L., Kanno, Y. & O’Shea, J. J. Proprotein convertase furin is preferentially expressed in T helper 1 cells and regulates interferon gamma. Blood 108, 983–985, doi:10.1182/blood-2005-09-3824 (2006).

10 Oksanen, A. et al. Proprotein convertase FURIN constrains Th2 differentiation and is critical for host resistance against Toxoplasma gondii. J Immunol 193, 5470–5479, doi:10.4049/jimmunol.1401629 (2014).

11 Huang, C. et al. Clinical features of patients infected with 2019 novel coronavirus in Wuhan, China. The Lancet 395, 497–506, doi:10.1016/S0140-6736(20)30183-5 (2020).

12 Chen, G. et al. Clinical and immunological features of severe and moderate coronavirus disease 2019. J Clin Invest 130, 2620–2629, doi:10.1172/JCI137244 (2020).

13 McGonagle, D., O’Donnell, J. S., Sharif, K., Emery, P. & Bridgewood, C. Immune mechanisms of pulmonary intravascular coagulopathy in COVID-19 pneumonia. The Lancet Rheumatology, doi:10.1016/S2665-9913(20)30121-1.

14 Diao, B. et al. Reduction and Functional Exhaustion of T Cells in Patients with Coronavirus Disease 2019 (COVID-19). medRxiv, 2020.2002.2018.20024364, doi:10.1101/2020.02.18.20024364 (2020).

15 Shimizu, A., Kondo, S., Sabe, H., Ishida, N. & Honjo, T. Structure and function of the interleukin 2 receptor: affinity conversion model. Immunol Rev 92, 103–120 (1986).

16 Liao, M. et al. Single-cell landscape of bronchoalveolar immune cells in patients with COVID-19. Nat Med, doi:10.1038/s41591-020-0901-9 (2020).

17 Bradley, A., Hashimoto, T. & Ono, M. Elucidating T Cell Activation-Dependent Mechanisms for Bifurcation of Regulatory and Effector T Cell Differentiation by Multidimensional and Single-Cell Analysis. Front Immunol 9, 1444, doi:10.3389/fimmu.2018.01444 (2018).

18 Ishikawa, E., Nakazawa, M., Yoshinari, M. & Minami, M. Role of Tumor Necrosis Factor-Related Apoptosis-Inducing Ligand in Immune Response to Influenza Virus Infection in Mice. Journal of Virology 79, 7658–7663, doi:10.1128/jvi.79.12.7658-7663.2005 (2005).

19 Ware, C. F. & Šedý, J. R. TNF Superfamily Networks: bidirectional and interference pathways of the herpesvirus entry mediator (TNFSF14). Current Opinion in Immunology 23, 627–631, doi:https://doi.org/10.1016/j.coi.2011.08.008 (2011).

20 Gonçalves-Carneiro, D., McKeating, J. A. & Bailey, D. The Measles Virus Receptor SLAMF1 Can Mediate Particle Endocytosis. Journal of Virology 91, e02255–02216, doi:10.1128/jvi.02255-16 (2017).

21 Fergusson, J. R. et al. CD161intCD8+ T cells: a novel population of highly functional, memory CD8+ T cells enriched within the gut. Mucosal Immunology 9, 401–413, doi:10.1038/mi.2015.69 (2016).

22 Bending, D. et al. A timer for analyzing temporally dynamic changes in transcription during differentiation in vivo. J Cell Biol, doi:10.1083/jcb.201711048 (2018).

23 Ono, M. & Tanaka, R. J. Controversies concerning thymus-derived regulatory T cells: fundamental issues and a new perspective. Immunol Cell Biol 94, 3–10, doi:10.1038/icb.2015.65 (2016).

24 Bailey-Bucktrout, S. L. et al. Self-antigen-driven activation induces instability of regulatory T cells during an inflammatory autoimmune response. Immunity 39, 949–962, doi:10.1016/j.immuni.2013.10.016 (2013).

25 Miyao, T. et al. Plasticity of Foxp3(+) T cells reflects promiscuous Foxp3 expression in conventional T cells but not reprogramming of regulatory T cells. Immunity 36, 262–275, doi:S1074-7613(12)00040-4 [pii] 10.1016/j.immuni.2011.12.012 (2012).

26 O’Malley, J. T. et al. Signal transducer and activator of transcription 4 limits the development of adaptive regulatory T cells. Immunology 127, 587–595, doi:10.1111/j.1365-2567.2008.03037.x (2009).

27 Zhou, L. et al. TGF-beta-induced Foxp3 inhibits T(H)17 cell differentiation by antagonizing RORgammat function. Nature 453, 236–240, doi:10.1038/nature06878 (2008).

28 Miyara, M. et al. Functional delineation and differentiation dynamics of human CD4+ T cells expressing the FoxP3 transcription factor. Immunity 30, 899–911, doi:S1074-7613(09)00202-7 [pii] 10.1016/j.immuni.2009.03.019 (2009).

29 Schönrich, G. & Raftery, M. J. The PD-1/PD-L1 Axis and Virus Infections: A Delicate Balance. Frontiers in Cellular and Infection Microbiology 9, doi:10.3389/fcimb.2019.00207 (2019).

30 Trinchieri, G. Interleukin-10 production by effector T cells: Th1 cells show self control. Journal of Experimental Medicine 204, 239–243, doi:10.1084/jem.20070104 (2007).

31 Roncarolo, M. G., Gregori, S., Bacchetta, R., Battaglia, M. & Gagliani, N. The Biology of T Regulatory Type 1 Cells and Their Therapeutic Application in Immune-Mediated Diseases. Immunity 49, 1004–1019, doi:10.1016/j.immuni.2018.12.001 (2018).

32 Brockmann, L. et al. IL-10 Receptor Signaling Is Essential for T_R_1 Cell Function In Vivo. The Journal of Immunology 198, 1130–1141, doi:10.4049/jimmunol.1601045 (2017).

33 Couper, K. N., Blount, D. G. & Riley, E. M. IL-10: The Master Regulator of Immunity to Infection. The Journal of Immunology 180, 5771–5777, doi:10.4049/jimmunol.180.9.5771 (2008).

34 Ono, M. Control of regulatory T-cell differentiation and function by T-cell receptor signalling and Foxp3 transcription factor complexes. Immunology 160, 24–37, doi:10.1111/imm.13178 (2020).

35 Heng, T. S., Painter, M. W. & Immunological Genome Project, C. The Immunological Genome Project: networks of gene expression in immune cells. Nat Immunol 9, 1091–1094, doi:10.1038/ni1008-1091 (2008).

36 LaMere, S. A., Thompson, R. C., Komori, H. K., Mark, A. & Salomon, D. R. Promoter H3K4 methylation dynamically reinforces activation-induced pathways in human CD4 T cells. Genes & Immunity 17, 283–297, doi:10.1038/gene.2016.19 (2016).

37 Ono, M. et al. Foxp3 controls regulatory T-cell function by interacting with AML1/Runx1. Nature 446, 685–689, doi:nature05673 [pii] 10.1038/nature05673 (2007).

38 Bending, D. et al. A temporally dynamic Foxp3 autoregulatory transcriptional circuit controls the effector Treg programme. EMBO J, doi:10.15252/embj.201899013 (2018).

39 Miyao, T. et al. Plasticity of Foxp3(+) T cells reflects promiscuous Foxp3 expression in conventional T cells but not reprogramming of regulatory T cells. Immunity 36, 262–275, doi:S1074-7613(12)00040-4 [pii] 10.1016/j.immuni.2011.12.012 (2012).

40 Bailey-Bucktrout, S. L. et al. Self-antigen-driven activation induces instability of regulatory T cells during an inflammatory autoimmune response. Immunity 39, 949–962, doi:10.1016/j.immuni.2013.10.016 (2013).

41 Wu, C. et al. Risk Factors Associated With Acute Respiratory Distress Syndrome and Death in Patients With Coronavirus Disease 2019 Pneumonia in Wuhan, China. JAMA Internal Medicine, doi:10.1001/jamainternmed.2020.0994 (2020).

42 Zheng, K. I. et al. Letter to the Editor: Obesity as a risk factor for greater severity of COVID-19 in patients with metabolic associated fatty liver disease. Metabolism 108, 154244–154244, doi:10.1016/j.metabol.2020.154244 (2020).

43 Meng, X. et al. Statins induce the accumulation of regulatory T cells in atherosclerotic plaque. Molecular medicine (Cambridge, Mass.) 18, 598–605, doi:10.2119/molmed.2011.00471 (2012).

44 George, J. et al. Regulatory T cells and IL-10 levels are reduced in patients with vulnerable coronary plaques. Atherosclerosis 222, 519–523, doi:10.1016/j.atherosclerosis.2012.03.016 (2012).

45 Brennan, A. M. & Mantzoros, C. S. Drug Insight: the role of leptin in human physiology and pathophysiology—emerging clinical applications. Nature Clinical Practice Endocrinology & Metabolism 2, 318–327, doi:10.1038/ncpendmet0196 (2006).

46 De Rosa, V. et al. A Key Role of Leptin in the Control of Regulatory T Cell Proliferation. Immunity 26, 241–255, doi:https://doi.org/10.1016/j.immuni.2007.01.011 (2007).

47 Visperas, A. & Vignali, D. A. A. Are Regulatory T Cells Defective in Type 1 Diabetes and Can We Fix Them? Journal of immunology (Baltimore, Md.: 1950) 197, 3762–3770, doi:10.4049/jimmunol.1601118 (2016).

48 Pearce, Erika L. & Pearce, Edward J. Metabolic Pathways in Immune Cell Activation and Quiescence. Immunity 38, 633–643, doi:10.1016/j.immuni.2013.04.005 (2013).

49 Gabriel, S. S. & Kallies, A. Sugars and fat – A healthy way to generate functional regulatory T cells. European Journal of Immunology 46, 2705–2709, doi:10.1002/eji.201646663 (2016).

50 Dang, E. V. et al. Control of T(H)17/T(reg) balance by hypoxia-inducible factor 1. Cell 146, 772–784, doi:10.1016/j.cell.2011.07.033 (2011).

51 Sugiura, D. et al. Restriction of PD-1 function by <em>cis</em>-PD-L1/CD80 interactions is required for optimal T cell responses. Science 364, 558–566, doi:10.1126/science.aav7062 (2019).

52 McNally, B., Ye, F., Willette, M. & Flaño, E. Local Blockade of Epithelial PDL-1 in the Airways Enhances T Cell Function and Viral Clearance during Influenza Virus Infection. Journal of Virology 87, 12916–12924, doi:10.1128/jvi.02423-13 (2013).

53 Stanciu, L. A. et al. Expression of Programmed Death–1 Ligand (PD-L) 1, PD-L2, B7-H3, and Inducible Costimulator Ligand on Human Respiratory Tract Epithelial Cells and Regulation by Respiratory Syncytial Virus and Type 1 and 2 Cytokines. The Journal of Infectious Diseases 193, 404–412, doi:10.1086/499275 (2006).

54 Pipkin, M. E. & Rao, A. SnapShot: effector and memory T cell differentiation. Cell 138, 606.e601–606.e6062, doi:10.1016/j.cell.2009.07.020 (2009).

55 Furmanski, A. L. et al. Tissue-Derived Hedgehog Proteins Modulate Th Differentiation and Disease. The Journal of Immunology 190, 2641–2649, doi:10.4049/jimmunol.1202541 (2013).

56 Kuczma, M. et al. Foxp3-Deficient Regulatory T Cells Do Not Revert into Conventional Effector CD4^+^ T Cells but Constitute a Unique Cell Subset. The Journal of Immunology 183, 3731–3741, doi:10.4049/jimmunol.0800601 (2009).

57 Stuart, T. et al. Comprehensive Integration of Single-Cell Data. Cell 177, 1888–1902.e1821, doi:10.1016/j.cell.2019.05.031 (2019).

58 Yu, G., Wang, L.-G., Han, Y. & He, Q.-Y. clusterProfiler: an R Package for Comparing Biological Themes Among Gene Clusters. OMICS: A Journal of Integrative Biology 16, 284–287, doi:10.1089/omi.2011.0118 (2012).

